# Linker histone H1 regulates defense priming and immunity in plants

**DOI:** 10.1101/2022.04.11.487821

**Authors:** Arsheed H. Sheikh, Kashif Nawaz, Naheed Tabassum, Marilia Trapp, Hanna Alhoraibi, Naganand Rayapuram, Heribert Hirt

## Abstract

Linker H1 histones play an important role in animal and human pathogenesis, but their function in plant immunity is poorly understood. Here, we analyzed mutants of the three canonical variants of Arabidopsis H1 histones, namely H1.1, H1.2 and H1.3. We observed that double *h1*.*1h1*.*2* and triple *h1*.*1h1*.*2h1*.*3* (*3h1*) mutants were resistant to *Pseudomonas syringae* and *Botrytis cinerea* infections. Transcriptome analysis of *3h1* mutant plants showed that histone H1s play a key role in regulating the expression of early and late defense genes upon pathogen challenge. Moreover, *3h1* mutant plants showed enhanced production of reactive oxygen species and activation of mitogen activated protein kinases upon pathogen-associated molecular pattern (PAMP) treatment. However, *3h1* mutant plants were insensitive to priming with flg22, a well-known bacterial PAMP (pathogen-associated molecular pattern) which induces enhanced resistance in WT plants. The defective defense response in *3h1* was correlated with the enhanced DNA methylation and reduced H3K56ac levels upon priming. Our data place H1 as a molecular gatekeeper in governing dynamic changes in the chromatin landscape of defense genes during plant pathogen interaction.

## Introduction

Plants face constant changes in their homeostasis, so they have evolved complex mechanisms to deal with external abiotic and biotic stresses. Plants can sense microbes by membrane-localized <u>p</u>attern recognition <u>r</u>eceptors (PRRs) of conserved <u>p</u>athogen/<u>m</u>icrobe-<u>a</u>ssociated <u>m</u>olecular <u>p</u>atterns (PAMPs/MAMPs), thereby triggering <u>P</u>AMP/<u>M</u>AMP triggered immunity (PTI/MTI) (Jones & Dangl, 2006). After recognition, downstream signaling responses are triggered, which include production of Reactive Oxygen Species (ROS), activation of <u>M</u>itogen-<u>A</u>ctivated <u>P</u>rotein <u>K</u>inases (MAPKs), and activation of defense genes (Bigeard *et al*, 2015). PTI also changes the plant defense hormone profiles (like salicylic acid and jasmonic acid) to optimize the immune response toward pathogens (Berens *et al*, 2017). Successful pathogens deliver effector proteins into the plant cells to overcome PTI, causing disease in susceptible plants called <u>e</u>ffector-triggered <u>s</u>usceptibility (ETS). However, resistant plants recognize effectors via intracellular <u>n</u>ucleotide-binding domain leucine-rich <u>r</u>epeat (NLR) receptors to mount <u>e</u>ffector triggered immunity (ETI) (Thomma *et al*, 2011).

In plants, a previous stressful experience modulates the cellular machinery to confer a robust response upon a recurring stress; a phenomenon called priming (Conrath *et al*, 2015). Priming of plants can efficiently increase the tolerance and survival of plants to different stresses. One of the well-characterized PAMPs is a 22-amino-acid long epitope of *Pseudomonas* flagellin (flg22), which is recognized by the FLS2 receptor (Chinchilla *et al*, 2007). Pretreatment with flg22 induces defense priming in Arabidopsis enhancing plant resistance to subsequent challenge with pathogenic *Pseudomonas* (Gong *et al*, 2019).

The execution of early and late immune responses upon pathogen recognition are fundamentally modulated by optimal gene expression (Birkenbihl *et al*, 2017; Winkelmüller *et al*, 2021). Chromatin changes and nucleosome dynamics fine-tune gene expression in stress conditions. Besides core nucleosomal histones, linker histones are the major constituents of eukaryotic nucleosomes (Kasinsky *et al*, 2001). In eukaryotes, linker histone H1 has a conserved tripartite structure consisting of 1) a short and flexible N-terminal tail, 2) a dyad binding central globular domain (GH1), and 3) a structurally disordered lysine-rich C-terminal tail (Zhou *et al*, 2013). Linker histone H1 binds to the nucleosome to facilitate chromatin folding usually into higher order structures (Fyodorov *et al*, 2018). In addition to regulating basic biological processes like DNA replication, chromosome segregation, and DNA repair, H1 regulates gene expression by modulating RNA polymerase II accessibility to chromatin (Hergeth & Schneider, 2015; Fan *et al*, 2005; Fyodorov *et al*, 2018).

H1 deposition is part of the crosstalk with the epigenetic landscape, notably DNA methylation and histone H3 methylation (Rupp & Becker, 2005; Yang *et al*, 2013; Rutowicz *et al*, 2019). H1 mutants are challenging to study owing to the high functional redundancy between H1 variants. Deletion of H1 in mouse and Drosophila is lethal but not in other organisms like Tetrahymena, yeast, fungi and worms (Shen *et al*, 1995; Fyodorov *et al*, 2018). Arabidopsis has three canonical variants of H1 namely, H1.1, H1.2, and stress-regulated H1.3 (Kotliński *et al*, 2017). Interestingly, the mutants are viable with morphological and developmental defects (Rutowicz *et al*, 2019). In Arabidopsis root and shoot tissues, H1.1 and H1.2 are ubiquitously expressed during vegetative development. At the genome level, H1 is distributed in heterochromatin and euchromatin showing enrichment at 3′ and 5′ ends of transposable elements (TEs). Over gene bodies, H1 presence is usually anticorrelated to transcription and H3K4me3 levels (Kotliński *et al*, 2017; Rutowicz *et al*, 2015). All DNA methylation patterns in Arabidopsis (i.e., CG, CHG, CHH), are regulated by H1 mainly in heterochromatin but also in euchromatin (Zemach *et al*, 2013; Rea *et al*, 2012). H1 possibly creates a structural barrier to hamper DDM1-mediated accessibility to DNA methyltransferases such as CMT2 (Zemach *et al*, 2013).

In this study, we demonstrate that Arabidopsis H1 knockout mutants have elevated basal immune levels and are resistant against a bacterial and a fungal pathogen. However, *3h1* mutant plants are compromised in flg22 triggered priming. The enhanced DNA methylation and reduced H3K56ac profiles in *3h1* after flg22-treatment provide strong evidence of the molecular mechanism of the priming deficient phenotype of *3h1* mutant.

## Results

### Linker histone H1 regulate plant immunity against pathogen infection

In an effort to understand the role of linker histone proteins in plant immunity, we analyzed Arabidopsis H1 mutants for developmental and pathogen phenotypes. The H1 mutants used in the study were either knockouts of single, double or all three isoforms of H1. Interestingly, all H1 mutants were viable and grew well with no visible morphological defect in pots (Fig 1A). We tested 4-week-old single mutants (*h1*.*2* and *h1*.*3*), double mutants (*h1*.*1h1*.*2, h1*.*1h1*.*3* and *h1*.*2h1*.*3*) and the triple mutant *h1*.*1h1*.*2h1*.*3* (called *3h1* from now onwards) against virulent *Pseudomonas syringae* pv. *tomato* DC3000 (*Pst* DC3000). We observed that *h1*.*1h1*.*2* and *3h1* are significantly resistant to *Pst* DC3000 infection while single *h1s*, double *h*.*1*.*1h1*.*3* and *h1*.*2h1*.*3* mutants allow proliferation of *Pst* DC3000 to a similar extent as found in wild type (WT) plants (Fig 1B). These observations suggest that two main isoforms H1.1 and H1.2 together are important in modulating plant immunity. As *3h1* showed the strongest resistance phenotype, we carried out the rest of the analysis using the triple *3h1* mutant. When challenged with the fungal pathogen *Botrytis cinerea, 3h1* plants efficiently restricted fungal infection compared with WT plants as depicted by the smaller lesion size (Fig 1C).

**Figure 1.**
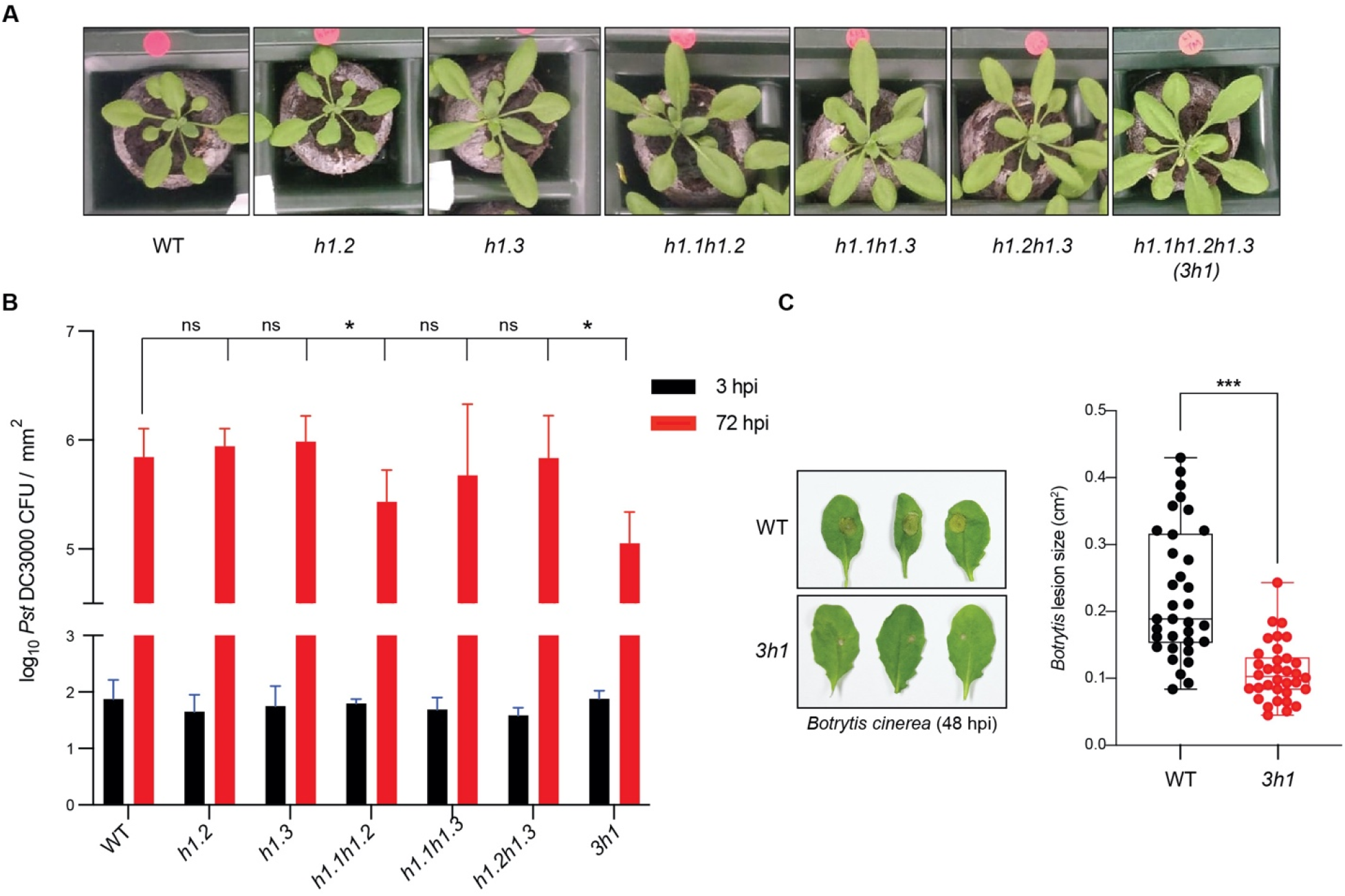
Linker histone mutant *3h1* is resistant to pathogen infection. **(A)** Morphology of 4-week-old wild-type (WT), single (*h1*.*2, h1*.*3*), double (*h1*.*1h1*.*2, h1*.*1h1*.*3, h1*.*2h1*.*3*) and triple (*h1*.*1h1*.*2h1*.*3* or *3h1*) mutants of Arabidopsis H1 grown on pots. **(B)** Quantification of bacterial colonies in different *h1* mutants after spray inoculation with *Pseudomonas syringae* pv DC3000 (*Pst* DC3000) at 3 and 72 hours post infection (hpi) **(C)** Quantification of fungal infection as reflected by the lesion size in the WT and *3h1* after 48 hours of *Botrytis cinerea* drop inoculation on 4 week-old plants. The data is represented as mean ± SEM where * indicate P < 0.05 compared to WT, as determined by ANOVA significance test.

### *3h1* mutant plants have elevated basal immune responses

To examine the effect of the loss of linker histones on basal immune responses, we performed a number of flg22-induced early and late PTI readouts in *3h1* mutant as compared to WT plants. For early PTI events, *3h1* plants produced higher levels of reactive oxygen species (ROS) than WT plants upon flg22 or chitin treatment (Fig 2A, EV1A). We also observed an elevated activation of the mitogen-activated protein kinases MPK3, 4 and 6 in *3h1* mutant as compared to WT after 15 and 30 min of flg22 elicitation in both adult plants and seedlings (Fig 2B, EV1B). The key plant enzymes producing ROS in responses to pathogen defense are RBOHD and RBOHF (Torres *et al*, 2002). We observed a higher level of both *RBOHD* and *RBOHF* levels in *Pst* DC3000 treated *3h1* plants (Fig 2C-D). Also, *PR1* defense gene expression was elevated in *3h1* mutant while a dynamic expression of *FRK1* and *MYB51* was observed before and after *Pst* DC3000 treatment (Fig 2E, EV1C, D). The defense related hormones like salicylic acid (SA) and jasmonic acid (JA) are key to optimize immune outputs, we thus quantified SA and JA in *3h1* mutant plants after pathogen infection. We observed elevated levels of SA in adult *3h1* plants compared to WT when challenged with *Pst* DC3000, while there was a subtle but insignificant increase in JA levels (Fig 2F, EV1E). Together, these results indicate that a deficiency in linker H1 affects basal immune responses in plants, possibly through H1-mediated regulation of the plant gene expression.

**Figure 2.**
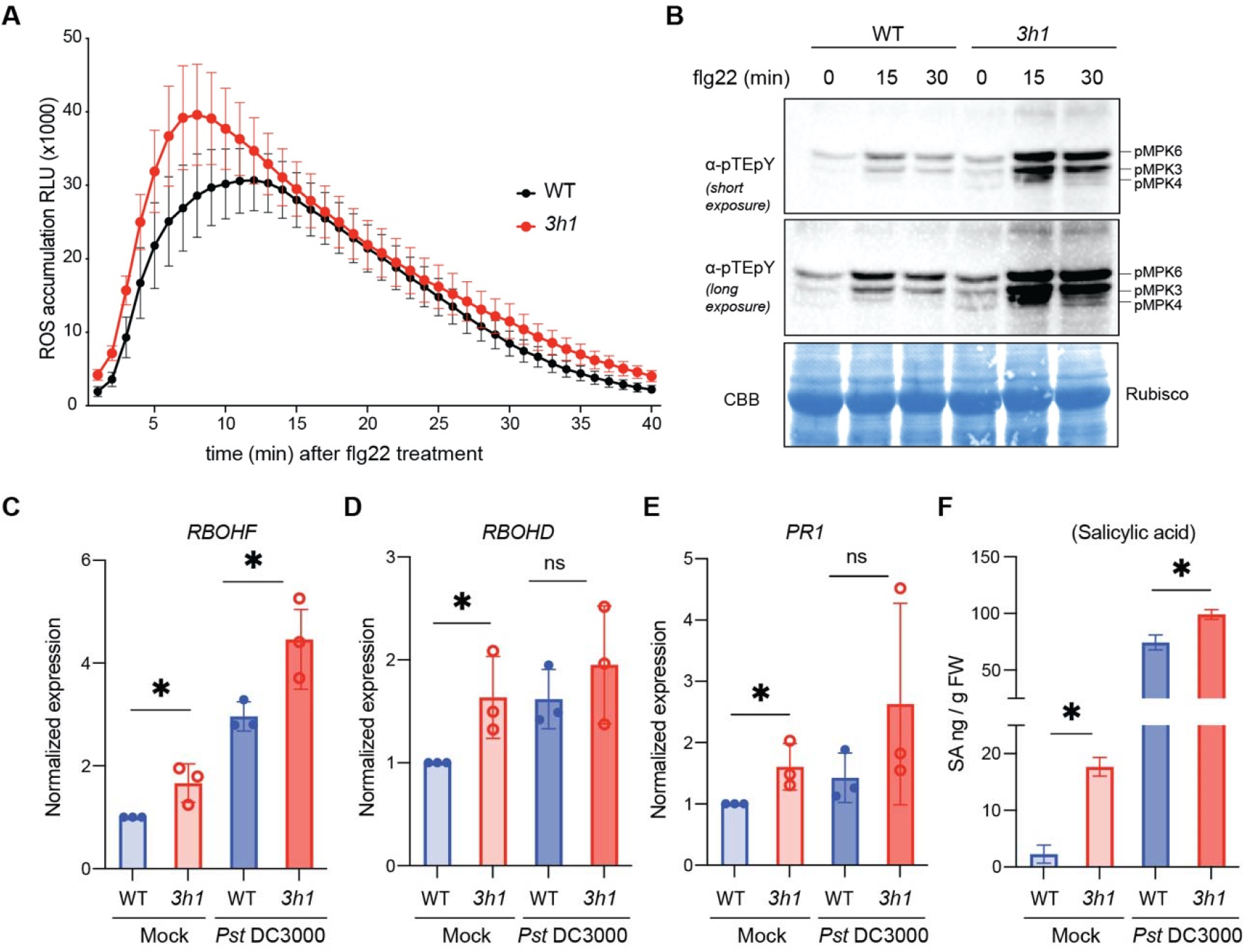
*3h1* mutant shows elevated innate immunity levels. **(A**) Reactive oxygen species (ROS) levels quantified in WT and *3h1* leaf discs triggered over 40 min by 1 μM flg22 treatment. **(B**) MAPK activation monitored with anti-pTEpY antibody as indicated with phosphorylation of MPK6, MPK3 and MPK4 in WT and *3h1* plants after 1 μM flg22 treatment for 15 and 30 min. CBB stain of Rubisco serves as a loading control. **(C, D, E)** Expression of key innate immunity genes *RBOHD, RBOHF* and *PR1* after 6h of *Pst* DC3000 infection. **(F)** Salicylic acid (SA) quantification in WT and *3h1* plants is shown as ng/g of fresh weight. The data are shown as means ± SEMs from three replicates. Asterisk indicates a significant difference with P < 0.05.

### H1 regulates the plant defense gene expression profile

In order to understand the immunity phenotype of the *3h1* mutant, we performed RNA sequencing of adult WT and *3h1* plants after challenge with *Pst* DC3000 for early (6h) and late (24h) defense responses (Fig 3A). Principal components analysis (PCA) revealed that biological replicates were clustered together (Fig EV2A), indicating good reproducibility of our experiments. Additionally, the 0 to 24 hpi samples were separated in the PCA plot, suggesting that gene expression changed massively as disease progressed (Fig EV2A). Differential gene expression analysis identified more than 1600 genes already affected by the genotype before challenge with *Pst* DC3000 (Fig EV2B). More than 1000 genes were upregulated, and around 600 genes were downregulated in *3h1* in control conditions (Fig EV2B). When compared to WT, *3h1* plants displayed significant differences in the abundance of 924 (638 up- and 286 downregulated) and 2151 (1145 up- and 1006 downregulated) genes after 6 and 24 hours of infection, respectively (Fig EV2B). Hierarchical clustering identified ten gene clusters with distinct changes in abundance in response to *Pst* DC3000 pathogen in *3h1* mutant plants compared to WT (Fig 3B). Notably, the transcripts in cluster 1, which were highly upregulated in WT at 24h, were repressed in *3h1* mutant plants upon *Pst* DC3000 treatment. This cluster contained a number of defense related genes, including *WRKY38* and *WRKY62*, which are both negative regulators of immunity (Kim *et al*, 2008). We confirmed the behavior of this cluster by qRT-PCR analysis of *WRKY38* and *WRKY62*, which both showed reduced expression in *3h1* post-infection by *Pst* DC3000 (Fig. 3C). The transcripts of cluster 5, including immunity-related defensins and cytochrome P450 family proteins, were highly upregulated in *3h1* but not in WT at 6h of infection (Fig 3B). Upon pathogen challenge, transcripts of cluster 6 and 7 genes showed highly elevated levels in *3h1* mutant plants at 24 h (Fig 3B). Whereas the GO terms of cluster 6 genes were associated with cell wall and polysaccharide metabolism, cluster 7 genes contained a number of chromatin related proteins, including the histone acetyltransferase HAC2 of the CBP family 2 and the histone deacetylase HDA18 of the RPD3/HDA1 superfamily (Fig 3B). By qRT-PCR, we observed that *HDA18* and *HAC2* levels strongly increased in *3h1* after *Pst* DC3000 infection compared to WT plants, eluding to the possible fine-tuning of gene expression by H1 (Fig 3D). Interestingly, cluster 9 genes, encoding flavonoid and anthocyanin secondary metabolite genes, were downregulated upon pathogen infection in WT but not in *3h1*, which showed elevated levels of these pathways before pathogen challenge. Together, these results suggest an essential role of H1 histones in orchestrating both early and late responsive plant defense genes.

**Figure 3.**
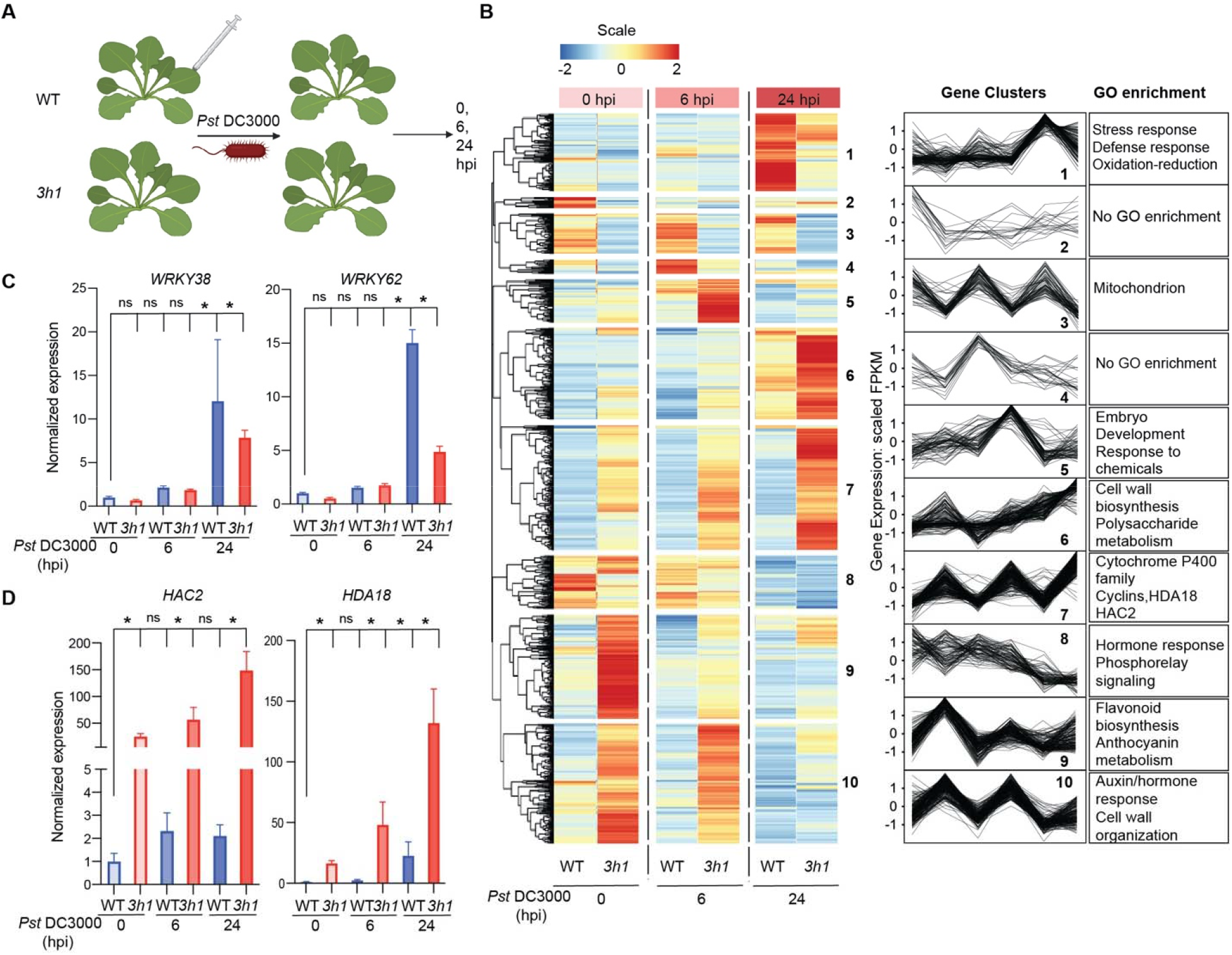
Dynamic transcriptome of *3h1* during pathogen infection **(A)** Schematic diagram of RNA seq procedure where 4 week-old adult plants were infected with *Pst* DC3000 and samples harvested at 0, 6 and 24 hours after infection. **(B)** Hierarchical clustering of *Pst* DC3000-infected (6 and 24 hours) WT and *3h1* plants compared to control based on differentially expressed RNA transcripts from mRNA seq using DESeq2 (2.0 FC, FDR padj< 0.01). Transcript fold-change depicted as color scale demonstrates log2 fold changes compared to mean for each transcript across genotype and condition. GO and pathway enrichment terms depicted for the gene clusters against the background of all transcripts in the heatmap. **(C, D)** qRT-PCR validation of selected genes which showed altered expression pattern between WT and *3h1* before and after infection. The data are shown as means ± SEMs from three replicates. Asterisk indicates a significant difference with P < 0.05 from 2way ANOVA.

### flg22 induced defense priming is attenuated in *3h1*

To understand whether H1 plays a role in defense priming, we pretreated WT and *3h1* plants with flg22 for 24 hours and performed *Pst* DC3000 pathogen infection 24 h later. Compared to water pretreatment, flg22 treated WT showed reduced disease symptoms of *Pst* DC3000 infection (Fig 4A, B). In contrast, *3h1* plants were insensitive to flg22-induced priming as shown by the similar bacterial titers in *3h1* mutant plants before and after flg22 treatment (Fig 4A, B). The reshaping of the transcriptome in pathogen-challenged tissues is one of the attributes of defense priming (Mauch-Mani *et al*, 2017). To understand the compromised defense priming phenotype in *3h1*, we performed transcriptome analysis of flg22 pre-treated WT and *3h1* plants challenged with *Pst* DC3000 at early (6h) and late (24h) time points (Fig 4C). PCA analysis showed the reliability of our biological replicates, and the Venn diagrams represent the numbers of up and downregulated genes after *Pst* DC3000 infection (Fig EV2C, D). Cluster 1 is comprised of 154 genes that get strongly upregulated in WT plants during early *Pst* DC3000 (6h) infection while the expression is not altered in *3h1* after infection (Fig 4D). Interestingly, GO term analysis shows strong enrichment of these genes in innate immunity and defense signaling. One of the key innate defense marker genes, *WRKY29* (Asai *et al*, 2002), was not induced in flg22-treated *3h1* plants after *Pst* DC3000 infection, suggesting a possible explanation for the compromised defense priming phenotype in *3h1* mutant plants (Fig 4E). We also tested the expression pattern of other defense-related genes like *WRKY31* and *MAPKKK15* in Cluster 1 (Fig 4F,G and Fig EV3A). Their expression behavior was similar to *WRKY29*, suggesting they might orchestrate the changes in plant defense in a concerted manner.

**Figure 4.**
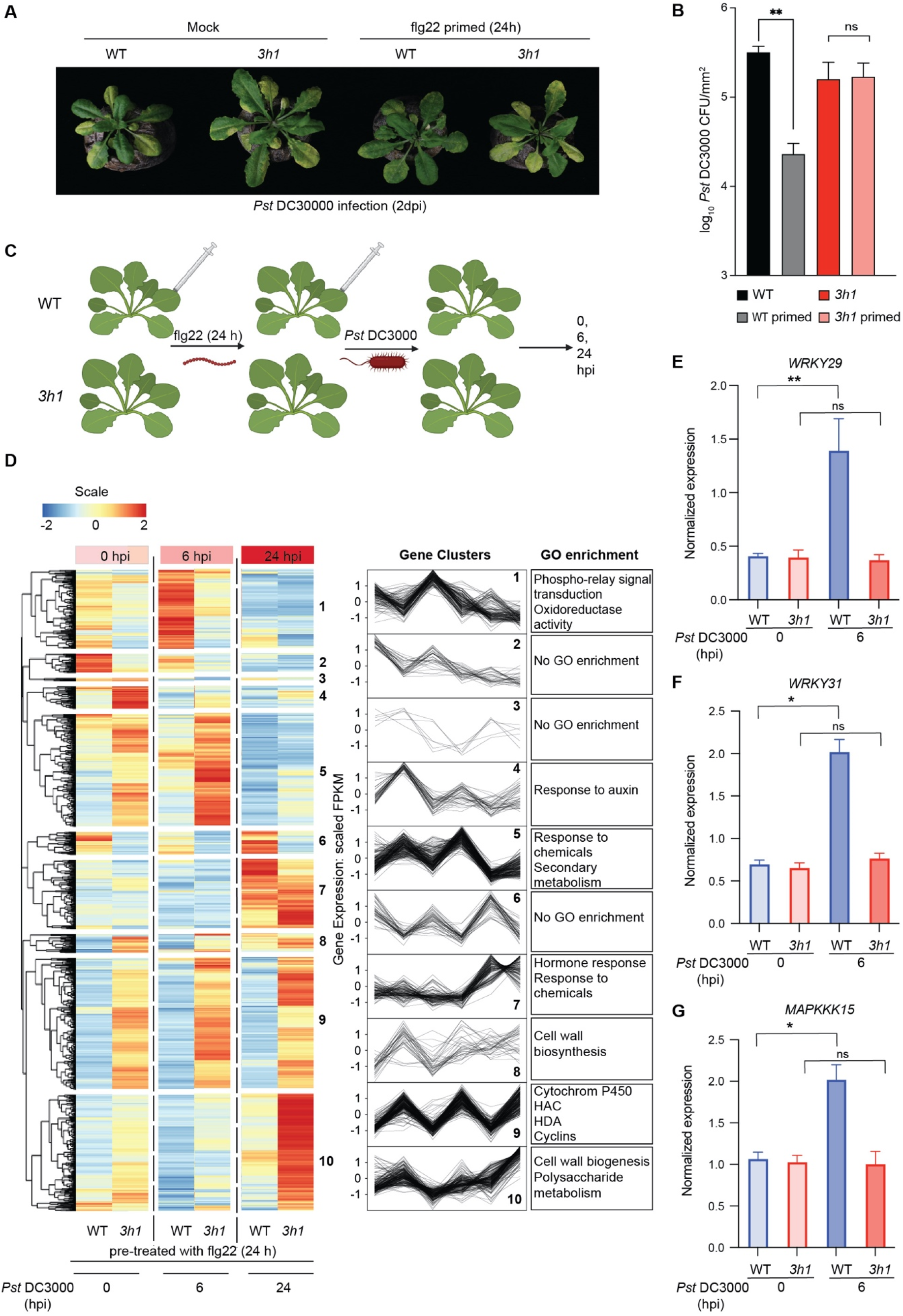
*3h1* is compromised in flg22-triggered defense priming **(A)** Altered disease symptom development in mock and flg22-treated WT and *3h1* adult plants. Photographs of representative infected plants were taken after 2 days of *Pst* DC3000 infection. **(B)** Bacterial growth of WT and *3h1* plants before and after 24h of flg22-treatemnt. **(C)** Schematic diagram of RNA seq where 4-week-old adult plants were first treated with 1 μM flg22 and after 24 h syringe infiltrated with *Pst* DC3000 and samples harvested at 0, 6 and 24 hours after infection. **(D)** Heat map showing differentially expressed transcripts of *Pst* DC3000-infected (6 and 24 hours) WT and *3h1* plants pre-treated 24 h prior with flg22. Transcript fold-change depicted as color scale demonstrates log2 fold changes compared to mean for each transcript across genotype and condition (FDR padj< 0.01). GO pathway enrichment terms depicted for the gene clusters against the background of all transcripts in the heatmap. **(E, F, G)** qRT-PCR validation of *WRKY29, WRKY31* and *MAPKKK15* genes which showed altered expression pattern between WT and *3h1* before and after infection when pre-treated with flg22. The data are shown as means ± SEMs from three replicates. Asterisk indicates a significant difference with P < 0.05 from multiple comparison 2way ANOVA.

Another stark difference was observed between *HAC2* and *HDA18* as their expression decreased after flg22 priming (Fig EV3B), which is in contrast to their expression in non-primed conditions (Fig 3D, E). These results suggest that an altered regulation of defense priming might occur at the epigenetic level in *3h1* mutant plants.

### *3h1* has elevated DNA methylation after flg22 treatment

The reshaping of the epigenetic landscape leads to a transiently enhanced local immunity followed by a poorly understood defense priming mechanism (Mauch-Mani *et al*, 2017). One of the key epigenetic modifications regulated by H1 histones is DNA methylation (Rutowicz *et al*, 2019; Bourguet *et al*, 2021). To further understand the defective priming in *3h1*, we analyzed genome-wide DNA methylation by whole-genome bisulfite sequencing (WGBS) in WT and *3h1* mutants pre-and post-flg22 treatment. Surprisingly, we observed a significant increase in overall methylation in *3h1* after flg22 treatment (Fig 5A, B). There were no significant changes in global methylation in WT plants after flg22 priming except for CHH sites (Fig 5B). H1 depletion resulted in an overall reduction in DNA methylation, especially in CHH and CHG sites for protein-coding regions (Fig 5B). Although the DNA methylation in all sequence contexts was increased in *3h1* over the TE-rich pericentromeric regions which contain the bulk of chromosome methylation, the average methylation over TEs in *3h1* was not massively altered after flg22 treatment as compared to protein coding regions. This indicates that flg22 treatment changes the methylation of <u>p</u>rotein <u>c</u>oding <u>g</u>enes (PCGs) in an H1-dependent manner (Fig 5C, D).

**Figure 5.**
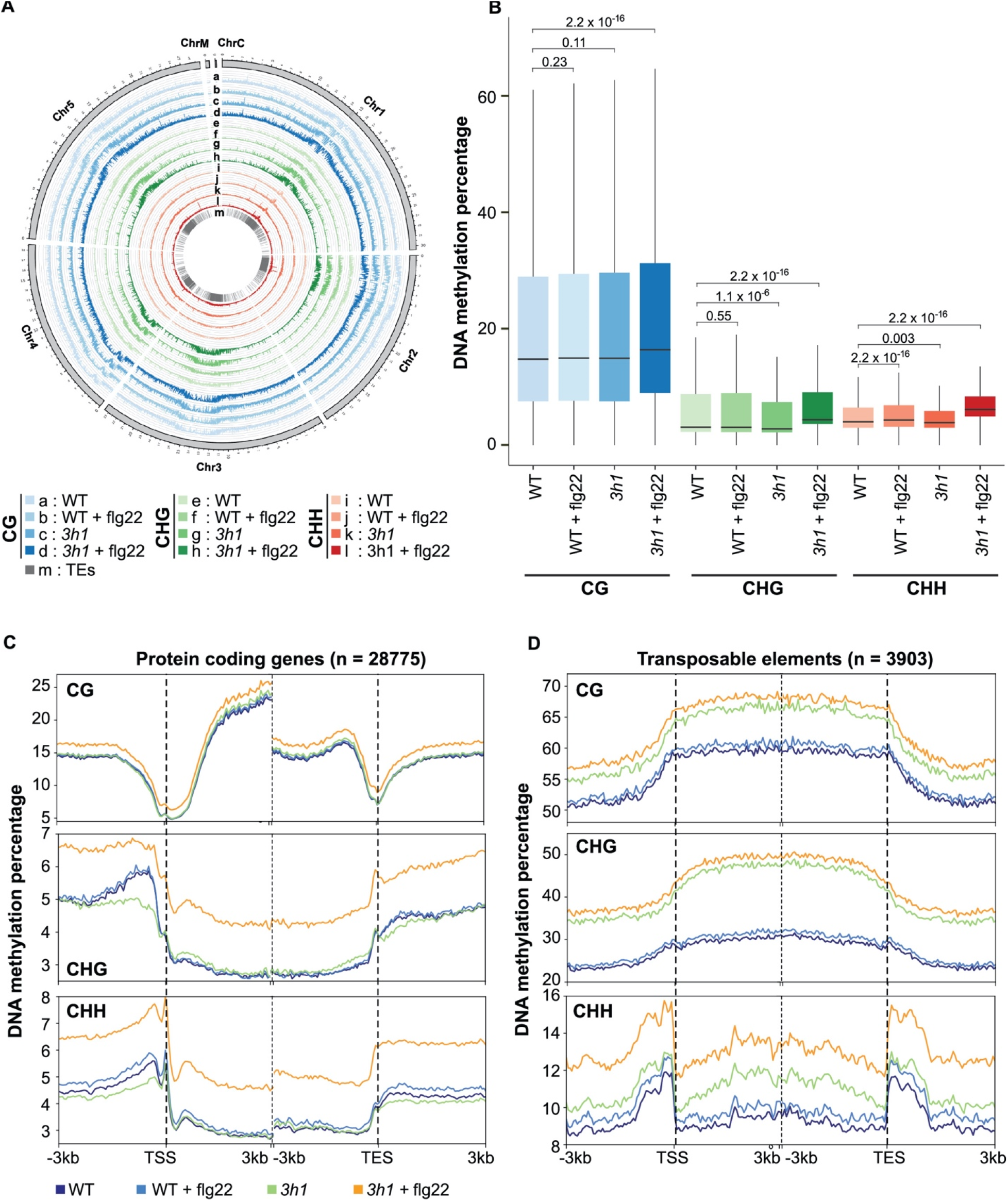
Global Methylation profile of *3h1* plants after flg22-treatment **(A)** Circos plot showing the genome wide methylation percentage tracks starting from outermost; CG WT, CG WT + flg22, CG *3h1*, CG *3h1* + flg22, CHG WT, CHG WT + flg22, CHG *3h1*, CHG *3h1* + flg22, CHH WT, CHH WT + flg22, CHH *3h1*, CHH *3h1* + flg22 and locations of TEs. **(B)** Boxplots of all methylation percentage and their significance with p-values. **(C, D)** Average plots of WT, WT + flg22, *3h1* and *3h1* + flg22 from all three contexts in protein coding genes (C) and TEs (D).

### flg22 triggers differential methylation in *3h1* plants

To understand the effects of elevated PCG methylation levels in flg22-treated *3h1* plants, we analyzed the gene expression and methylation profile changes in *3h1* plants compared to WT. Flg22 triggers differential regulation of 2433 genes in WT, of which 1509 are up- and 924 down-regulated, whereas flg22 triggers almost twice the number of DEGs in *3h1* (5160 genes, of which 2560 are up- and 2600 are down-regulated). This reflects that H1 histones suppress gene expression under normal circumstances, which is consistent with 1008 up- and 609 down-regulated genes in *3h1* mutant plants under control conditions compared to WT (Fig 6A). Interestingly in *3h1*, the number of flg22-induced DEGs is decreased and only 389 genes are up- and 154 down-regulated in flg22 treated *3h1* plants compared to flg22 treated WT. This decrease in DEGs could be a direct effect of increased global PCG methylation in *3h1* after flg22 treatment. To understand the direct relation of gene expression with DNA methylation, we divided the up- and down-regulated genes into either methylated or unmethylated loci. We observed that demethylation was necessary for changing gene expression, whereas methylation did not change for genes with an unchanged expression before or after flg22 treatment (Fig 6B). This is in agreement with the general assumption that demethylation is necessary to induce gene expression. One of the prominent gene clusters (Cluster 1, Fig 4D) where the gene expression massively increases at 6h of *Pst* DC3000 infection in flg22-treated WT plants but not in *3h1*, showed a stark opposite profile for both promoter and gene body methylation (Fig EV4A, B). Moreover, the 30 genes directly involved in plant innate defense showed a striking correlation between promoter and gene body methylation and repressed expression patterns in *3h1* as compared to WT (Fig 6C, EV4C). This further reiterates that H1 regulates flg22-induced methylation and subsequent gene expression changes in plants. As flg22 enhanced PCG DNA methylation in all contexts (CG, CHH, and CHG) in *3h1* mutant plants, we speculate that the altered gene expression of *WRKY29, HAC2*, and *HDA18* (Fig 4D, EV3B) might be due to enhanced methylation at these gene loci. As expected, we observed increased methylation, especially in the CHH context at these gene loci (Fig 6D).

**Figure 6.**
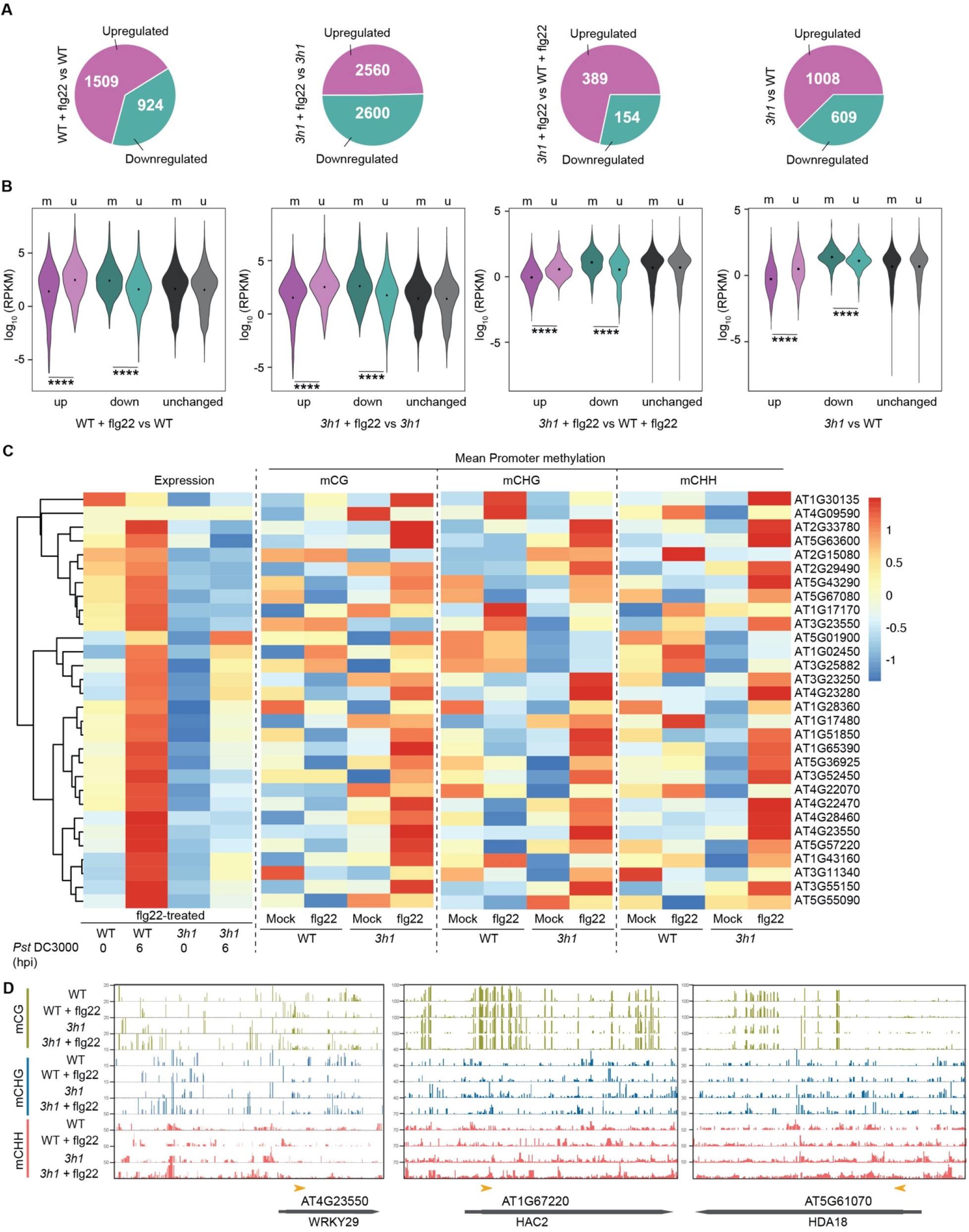
flg22 triggers differential methylation and gene expression in *3h1* **(A)** The number of differentially expressed genes triggered by flg22-treatment in WT and 3h1. **(B)** Violin plots showing the breakdown of mis-regulated transcripts in *3h1* (mutant versus WT) into methylated (m), unmethylated (u) either with or without flg22-treatment (****p < 0.0001, Wilcoxon test). **(C)** Heatmap of mean promoter methylation in all contexts (CG, CHG and CHH) on selected transcripts (30 genes) from Cluster 1 (154 genes) in flg22-treated WT and *3h1* samples which showed differential expression after *Pst* DC3000 infection (Cluster 1 of Fig 4D). **(D)** Genome Browser snapshots showing the distribution of CH, CHG and CHH methylation in representative genes *WRKY29, HAC2* and *HDA18*.

### H1 histones control H3K56ac levels in plants

H1 histones control the epigenetic landscape in animals by either promoting or inhibiting various H3 histone modifications (Willcockson *et al*, 2020). Since we also observed dynamic differential regulation of *HAC2* and *HDA18* in *3h1*, we speculated that H1 might also regulate the histone epigenetic profile. To this end, we tested the global levels of two well-known active and repressive marks H3K4m3 and H3K27me3, respectively, in flg22-primed and non-primed plants after *Pst* DC3000 infection. We observed no significant changes in these histone marks between *3h1* and WT plants (Fig EV5). Since H1 and H3K56ac can act as antagonistic regulators of nucleosome dynamics (Bernier *et al*, 2015), we also analyzed the levels of H3K56ac in *3h1* plants. Although H3K56ac levels were enhanced in *3h1* plants, we observed a dramatic decrease in H3K56ac levels in flg22-treated *3h1* mutant plants (Fig 7A). In contrast, flg22 treatment increased H3K56ac in WT plants, suggesting that H3K56ac is a crucial player in flg22-triggered priming in plants. Interestingly, the H3K56ac levels decreased upon *Pst* DC3000 infection, suggesting a possible strategy of the pathogen to overcome plant defense. The combined regulation of DNA methylation and histone modifications by H1 leads to a dynamic chromatin landscape (Yang *et al*, 2013). Finally, to test the state of chromatin compaction of some of the selected differentially regulated genes in *3h1*, we performed DNase I accessibility assays (Shu *et al*, 2013) (Fig 7B). We observed that WRKY29, *HAC2* and *WRKY38* were more accessible to DNase I digestion in *3h1*, suggesting an open chromatin state for these genes (Fig 7C). However, the accessibility of *HDA18* and *WRKY62* was not significantly changed in *3h1*, reflecting a more complex regulation of chromatin accessibility (Fig EV6A). Conversely, the overall accessibility of *HAC2, WRKY29* and *WRKY38* to some extent was restricted by flg22 treatment in *3h1 (*Fig EV6B, C). Taken together, our data suggest that a multifaceted regulation of the chromatin landscape by H1 histones via dynamic DNA methylation and H3K56 acetylation mainly accounts for the defective priming phenotype in *3h1* plants (Fig 7D).

**Figure 7.**
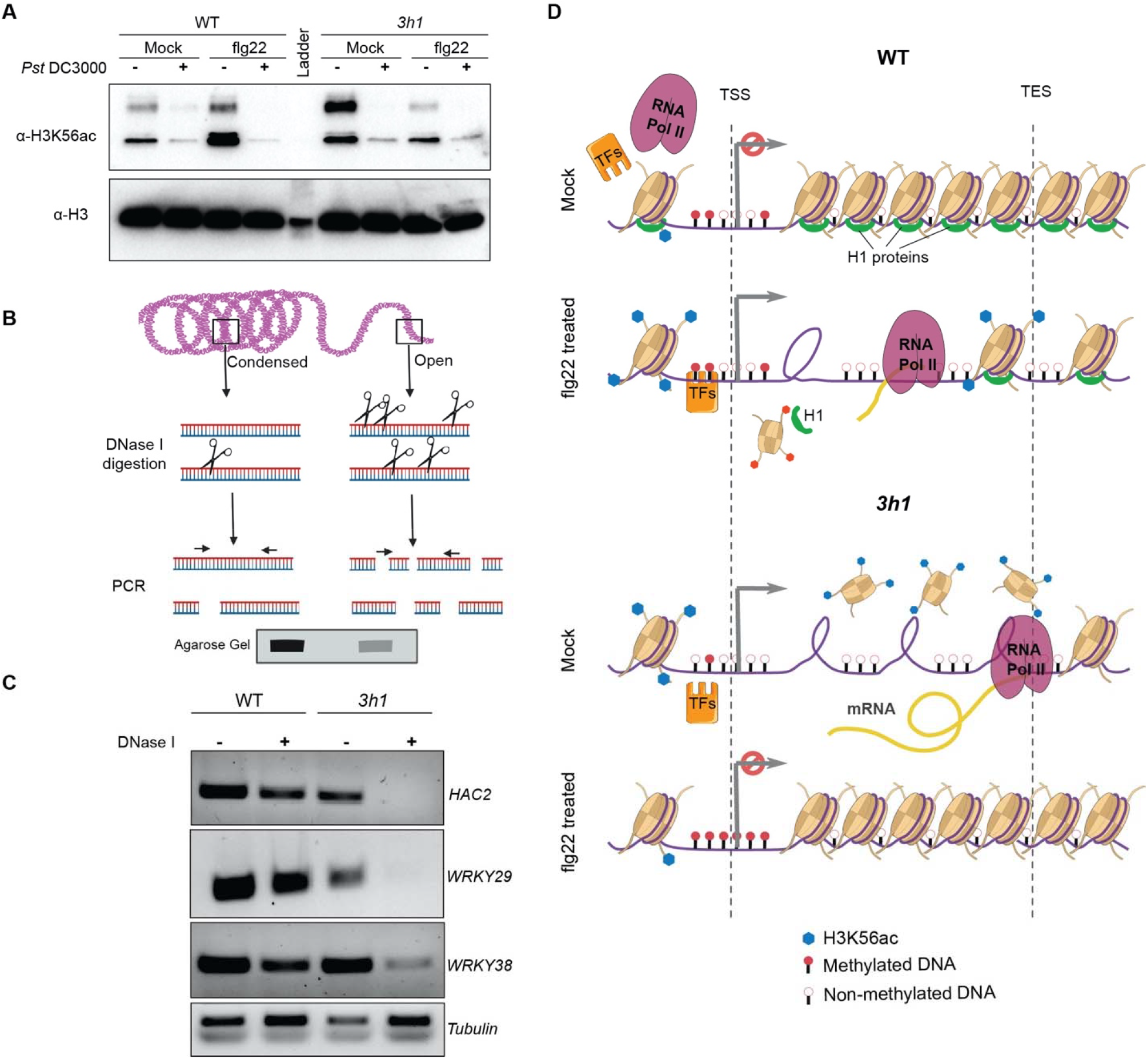
H1 controls the H3K56ac levels in plants **(A)** Western blot showing the H3K56ac levels in WT and *3h1* plants after water (mock) or flg22 treatment challenged with *Pst* DC3000 (6h). **(B)** Schematic representation showing the principle of the chromatin accessibility assay by DNase I digestion using PCR. Open chromatin is more frequently cut by limited DNase I digestion in small fragments giving reduced PCR signals while condensed chromatin which is less frequently cut gives strong PCR band signal. **(C)** DNase I accessibility PCR of *HAC2, WRKY29* and *WRKY38* in WT as compared to *3h1*. **(D)** Model showing the dynamic changes in epigenetic landscape in *3h1* after flg22 treatment as compared to WT Arabidopsis plants.

## Discussion

The role of linker histones is extensively studied in animal disease pathogenesis and progression (Ye *et al*, 2017). However, we lack an understanding of the role of H1 histones in plant immunity and disease. In this study, we aimed to understand the role of linker histone H1 in plant defense. The lack of developmental or pathogen phenotypes in single mutants of Arabidopsis H1.1 and H1.2 reflects the redundancy of the linker histones (Fig 1). In contrast, animal studies showed that single H1 mutations lead to severe developmental defects or diseases (Behrends & Engmann, 2020). This can partly be explained by the reduced number of H1 variants present in animals, while three isoforms of H1 exist in Arabidopsis (Kotliński *et al*, 2017). Accordingly, double *h1*.*1h1*.*2* and triple mutant *h1*.*1h1*.*2h*.*1*.*3* (*3h1*) plants show altered resistance to both bacterial and fungal pathogens. The developmental defects like stomatal spacing in *3h1* do not seem to influence the bacterial load in plants, as reflected by almost equal bacterial counts at 3 hpi (Fig 1B) (Rutowicz *et al*, 2019). In *3h1*, the enhanced disease resistance is associated with enhanced PTI induction as reflected by elevated ROS and MAPK activation (Fig 2A-B). The enhanced PTI marker gene expression and elevated defense hormone SA levels further support the resistance phenotype of *3h1* mutant plants. The transcriptome analysis also hints toward higher levels of secondary metabolites like phytoalexins and anthocyanins in *3h1* mutant plants in control conditions (Cluster 6,7 and 9, Fig 3B). These metabolites are known to regulate plant immunity against both bacterial and fungal pathogens (Piasecka *et al*, 2015) and could explain why *3h1* plants are resistant to both bacterial and fungal pathogens (Fig 1). Interestingly, after flg22 priming, WT plants show enhanced expression of defense-related genes like MAPKs and WRKYs compared to *3h1* (Cluster 1, Fig 4D) (Asai *et al*, 2002). The inability of flg22-treated *3h1* plants to mount a robust defense response after pathogen infection could lead to the observed non-priming phenotype (Fig 4A-B). The other major factors which explain the priming-deficient phenotype in *3h1* are the DNA methylation and histone acetylation changes in the mutant after flg22 treatment. The presence of H1 histones might present a physical barrier to PCG targeting DNA methyltransferases which are activated by flg22 priming. The interdependence of various methylation pathways might explain the increase in PCG methylation in all contexts in *3h1* after flg22 treatment (Du *et al*, 2015). However, the drop in CHG and CHH but not in CG PCG methylation at the promoters of genes in *3h1* mutant plants indicates a distinct regulation of these methylation patterns (Fig 5C). In contrast, the higher methylation of TEs in all contexts in *3h1* mutant plants is in agreement with the methylation patterns observed for long TEs in the double mutant *h1*.*1h1*.*2* (Fig 5D) (Zemach *et al*, 2013). The minor changes in TE methylation after flg22 treatment suggest that H1 histones primarily change the flg22-induced methylation in euchromatic genes (Fig 5D). Linker H1 histones are normally associated with the compaction of nucleosomes and hinder the access of the transcription machinery to genes (Fyodorov *et al*, 2018). This is also suggested by our data as we observe a higher number of differentially regulated genes in *3h1* under control conditions (Fig 5A). In the absence of H1, chromatin seems to be more accessible to enzymes for euchromatic histone modifications and antagonizes DNA methylation (Zemach *et al*, 2013). For example, the histone acetyltransferase INCREASED DNA METHYLATION (IDM1) is important for preventing DNA hypermethylation, while the histone deacetylase HDA6 affects DNA methylation particularly in rDNA (Qian *et al*, 2012; Probst *et al*, 2004). We also observed that the histone acetyltransferase *HAC2* and the deacetylase *HDA18* were strongly upregulated in *3h1* mutant plants, which could antagonize DNA methylation (Fig 3D). This H1 dependent interplay between DNA methylation and histone modifiers prompted us to analyze the levels of histone marks in *3h1* before and after flg22 treatment. The higher levels of the H3K56ac activation mark in the absence of H1 (*3h1*) suggest the greater access of transcription factors to the gene promoters (Fig 7A) (Bernier *et al*, 2015). This makes sense as H3K56 is close to the exit-entry points of the nucleosomal DNA superhelix which coincides with the position of linker histone H1. However, the minor changes observed in H3K4me3 and H3K27me3 in *3h1* compared to WT could be due to the cell type-specific and developmental stage-specific nature of these modifications (Geeven *et al*, 2015) (Fig EV5). Paradoxically, flg22 treatment downregulates *HAC2* and *HDA18* expression in *3h1* mutant plants, possibly leading to reduced histone H3K56ac levels, which may allow the DNA methylation machinery to increase PCG methylation. The intricate crosstalk between DNA methylation and histone acetylation in *3h1* plants before and after flg22 treatment can lead to structurally more dynamic chromatin accessibility to the transcription machinery. Taken together, we provide evidence for the role of H1 in governing dynamic changes in DNA methylation and histone acetylation during plant immunity.

## Materials and Methods

### Plant material and growth conditions

The experiments were performed by using Arabidopsis (*Arabidopsis thaliana*) lines in the Col-0 ecotype background unless stated otherwise. *h1*.*1*(SALK_N628430) and *h1*.*2* (GK-116E080) mutants were obtained from the European Arabidopsis Stock Center. *h1*.*3* (GT18298) in the L*er* background was obtained from the Cold Spring Harbor Laboratory collection. The *h1*.*1h1*.*2h1*.*3* (*3h1*) triple mutant was obtained by crossing *h1*.*1h1*.*2* with *h1*.*3* (Rutowicz *et al*, 2019). Plants were grown on soil (jiffy pots, http://www.jiffypot.com/), in plant growth chambers (Percival Scientific) under short-day conditions (8h light/ 16 h dark) at 22°C.

### Pathogen infection assays

*P. syringae tomato* pv. DC3000 (*Pst* DC3000) strain was grown on King B agar plates with 50 μg/ml rifampicin at 28°C. Bacteria were adjusted to 10^6^ cfu/ml in 10 mM MgCl_2_ and inoculated to four-week-old plants by either syringe or spray infection. Bacteria were released from three leaf discs (4mm) in 500 μl of 10 mM MgCl_2_ with 0.01% Silwet 77 by incubating at 28°C at 1000 rpm for 1 h. Serial dilutions were plated on LB medium with appropriated antibiotic. After incubation at 28°C, bacterial colonies were counted. The experiment was repeated three times with similar results.

*Botrytis cinerea* infection was performed as previously described (Rayapuram *et al*, 2021). Briefly, 4-week-old plants were inoculated with *Botrytis cinerea* (strain: B05.10) by placing a 5 μL droplet of a spore suspension (5×10^5^ spores/mL) on each rosette leaf (three fully expanded leaves per plant). Trays were covered by a transparent plastic lid to maintain high humidity. Lesion diameter was measured after 2 days of infection using ImageJ analysis tool.

### ROS Burst assay

ROS burst was determined by the luminol-based assay described before with modifications (Ranf *et al*, 2011). Leaf discs were incubated overnight in a white 96-well plate in water to reduce the wounding effect. Next day, 100 μl of reaction solution containing 50 μM of luminol (Sigma) and 10 μg/ml of horseradish peroxidase (Sigma) supplemented with 1 μM of flg22. The measurement was conducted immediately with the luminometer (GloMax, Promega), for a period of 40 min with a 1 min reading interval between readings. The measurements are indicated as means of RLU (Relative Light Units). The experiments were repeated three times with similar results.

### MAPK activation assays

Adult plants or seedlings treated with 1 μM of flg22 were harvested at indicated time points. Proteins were extracted using extraction buffer. The frozen seedlings were homogenized in 100 μL of extraction buffer (150 mM Tris-HCl, pH 7.5, 150 mM NaCl, 5 mM EDTA, 2 mM EGTA, 5% glycerol, 10 mM DTT and 1 x Pierce Protease and Phosphatase tablet (thermo#A32959). Twenty micrograms of protein were separated on a 10% polyacrylamide gel and transferred on a PVDF membrane. After overnight blocking with TBST-5% milk, the membrane was washed 3 times with TBST. Immunoblot analysis was performed using anti-phospho-p44/42 MAPK (1:5,000; Cell Signaling Technology) for 2 h as primary antibody and peroxidase-conjugated goat anti-rabbit antibody (1:10,000; Promega) for 1 h with 5 times 10 min washes in-between the incubations. The experiment was repeated twice with similar results.

### Hormone measurement

The extraction of phytohormones was performed as already described (Trapp *et al*, 2014). The compounds were quantified by HPLC-ESI-SRM, in a Thermo Fisher TQS-Altis Triple Quadrupole Mass Spectrometer coupled to a Thermo Scientific Vanquish MD HPLC system. The chromatographic separation was carried out in a UPLC column (Agilent Eclipse Plus C18, RRHD, 1.8 um, 2.1 × 50mm), and the compounds were eluted using water (A) and acetonitrile (B) as mobile phase at 0.6 mL/min and in a gradient elution mode as following: 10% B for 0.5 min, 10-55 % of B at 4.5min, 55-100% B at 4.7 min, 100% until 6.0 min, 100-10% B at 6.1% and 10% until 8 min. The column was kept at 55°C. The statistical significance from three replicates was evaluated by ANOVA followed by Tukey Test. (Tukey HSD).

### RNA Extraction and real-time quantitative PCR analysis

Total RNAs were extracted from 4-week-old adult plants using NucleoSpin RNA Plant (MACHEREY-NAGEL) kit, according to the manufacturer’s instructions. First strand cDNA was synthesised from 1μg of total RNAs using SuperScript First-Strand Synthesis System for RT-PCR according to the manual instructions. Then 2 μl of 10 times diluted cDNA was used for CFX96 Touch Real-Time PCR Detection System (Bio-Rad) for Syber green based quantitative PCR analysis. The specificity of amplification products was determined by melting curves. Tubulin was used as internal control for signals normalisation. The relative expression level of the selected genes was calculated based on ΔΔCt method. The primers used are listed in Table EV1

### RNA sequencing and Analysis

RNA seq was performed from mRNA libraries with 1µg of total plant RNA using TrueSeq standard mRNA Library Prep kit (Illumina). Pooled libraries were sequenced using Illumina HiSeq 4000 platform. RNA-seq generated 1538 million raw read pairs from 36 libraries (include three time points; 0hr, 6hr and 24h in methylation data). Adapters, primers, and low-quality bases were removed from the ends of raw reads using Trimmomatic v0.38 (Bolger *et al*, 2014). The resulting trimmed reads were mapped to the TAIR10 genome using Tophat2 v2.1.1 (Kim *et al*, 2013). Read counts were generated for all samples from corresponding bam files using BEDTools v2.29.0 (Quinlan & Hall, 2010). DESeq2 (Love *et al*, 2014) was run with read counts to perform DEGs between several conditions (comparison description: refer to RNA-seq excel file) with FDR ≤ 0.01.

Functional enrichment of DEGs was carried out with AgriGO (Tian *et al*, 2017) using default settings. GO terms with P ≤ 0.05 were considered significant, and the occurrence of at least five times in the background set was additionally required for DEGs.

### Whole Genome Bisulfite Seq and Analysis

Sequencing of the 12 libraries (4 conditions, 3 biological replicates each) resulted in 991 million read pairs of the Illumina HiSeq 4000 platform. Adapters were trimmed from the raw sequences using Trimmomatic v0.38. Subsequently, trimmed reads were mapped to the TAIR10 genome using Bowtie2 v2.2.5 (Langmead & Salzberg, 2012), and methylation calls were performed using Bismark v0.22.3 (Krueger & Andrews, 2011). Three filters were used to reduce false positives. First, for each position with k methylated reads mapping to it, the probability of it occurring through sequencing error (that is, unmethylated position appearing as methylated) was modeled using a binomial distribution B (n, p), where n is the coverage (methylated + unmethylated reads) and p is the probability of sequencing error (set to 0.01). We kept positions with k methylated reads if P (X≥ k) < 0.05 (post-FDR correction). Second, retained methylated positions had to have ≥ 1methylated read in all three biological replicates of at least one growth condition. Finally, the median coverage of retained positions across all 12 samples had to be ≥10.

### Assignment of genomic context to methylated cytosines

On the basis of the gene annotation of the TAIR10 (GFF3 file) and the positional coordinates of the methylated cytosines produced by Bismark, we annotated every methylated cytosine based on the genomic context, including whether the methylated position resides in a genic or intergenic region, and the distances to the 5′ and 3′ ends of each genomic feature (gene/intergenic region/exon/intron).

### PCA and correlation matrices

Median methylation levels of methylated genes and log2FPKM of expressed genes were shifted to be zero-centered and analyzed by PCA using the prcomp function in R. Using the same data, we calculated correlation matrices (Pearson correlation coefficient) and clustered samples with hclust implemented in R using complete linkage and Euclidean distance.

### Western Blotting of Histones

Nuclei were isolated from ground powder as already described (Ramirez-Prado *et al*, 2021). 5x SDS Loading dye was directly added to the nuclei and boiled at 85°C for 10 min and later loaded on 15% SDS-PAGE gel. Later Western blotting was performed as described in MAPK activation assays section.

### DNaseI Accessibility PCR

DNaseI (Promega) treatment was given to isolated nuclei for 5 min at 37°C and reaction was stopped by adding EDTA to the tubes. Later DNA was isolated from the samples and accessibility PCR assay performed as already described (Shu *et al*, 2013).

## Expanded View Figs

**Figure EV1.**
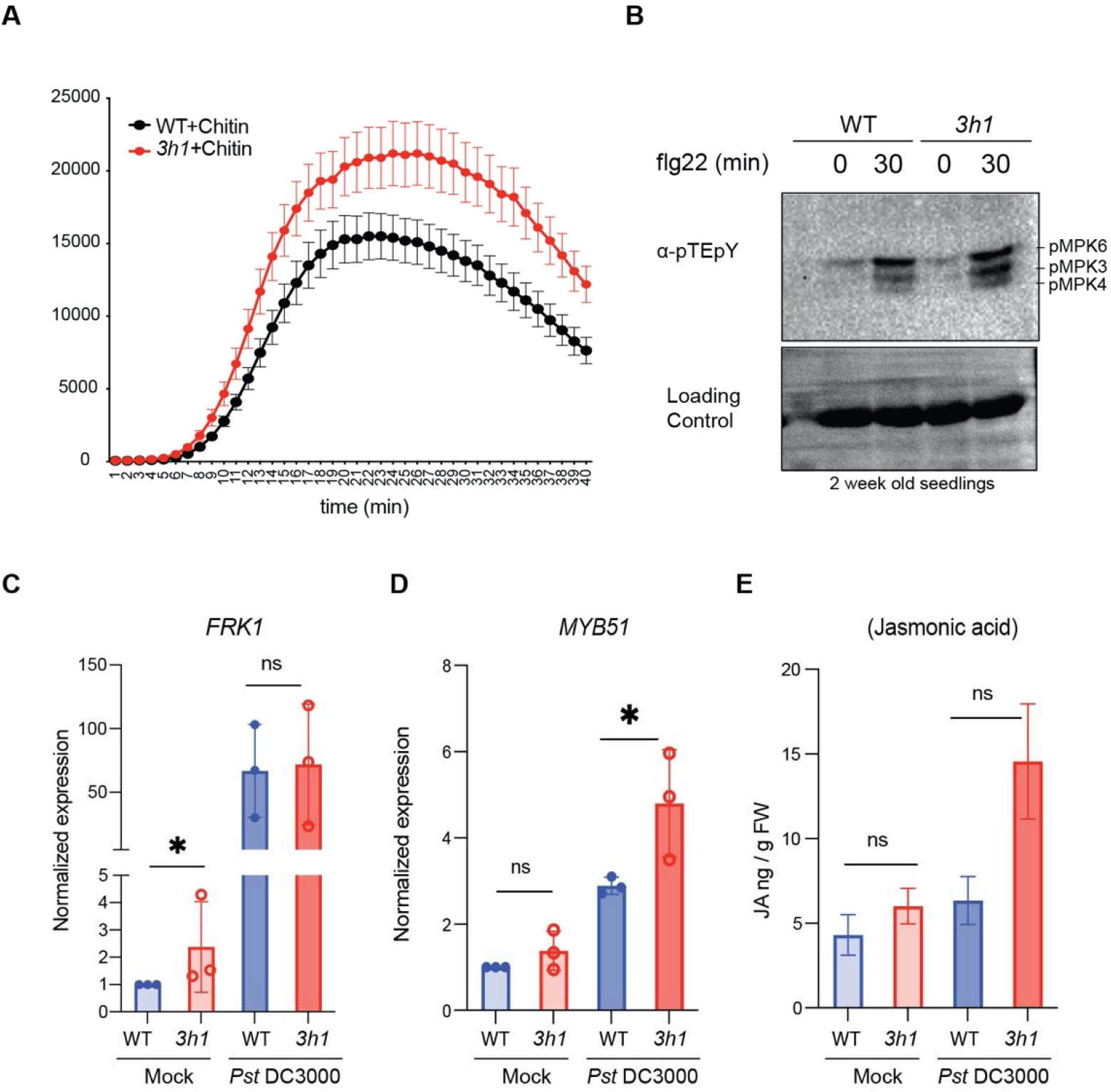
Innate immunity (PTI) levels in *3h1* **(A)** Reactive oxygen species (ROS) levels quantified in WT and *3h1* leaf discs triggered over 40 min by chitin treatment. **(B)** MAPK activation shown by anti-pTEpY western blot indicates the phosphorylation of MPK6, MPK3 and MPK4 in 2-week-old WT and *3h1* seedlings after 1 μM flg22 treatment for 30 min. CBB stain of Rubisco serves as a loading control. **(C, D)** Expression of innate immunity genes *FRK1* and *MYB51* after 6h of *Pst* DC3000 infection. **(E)** Jasmonic acid (JA) quantification in WT and *3h1* plants is shown as ng/g of fresh weight. The data are shown as means ± SEMs from three replicates. Asterisk indicates a significant difference with P < 0.05.

**Figure EV2.**
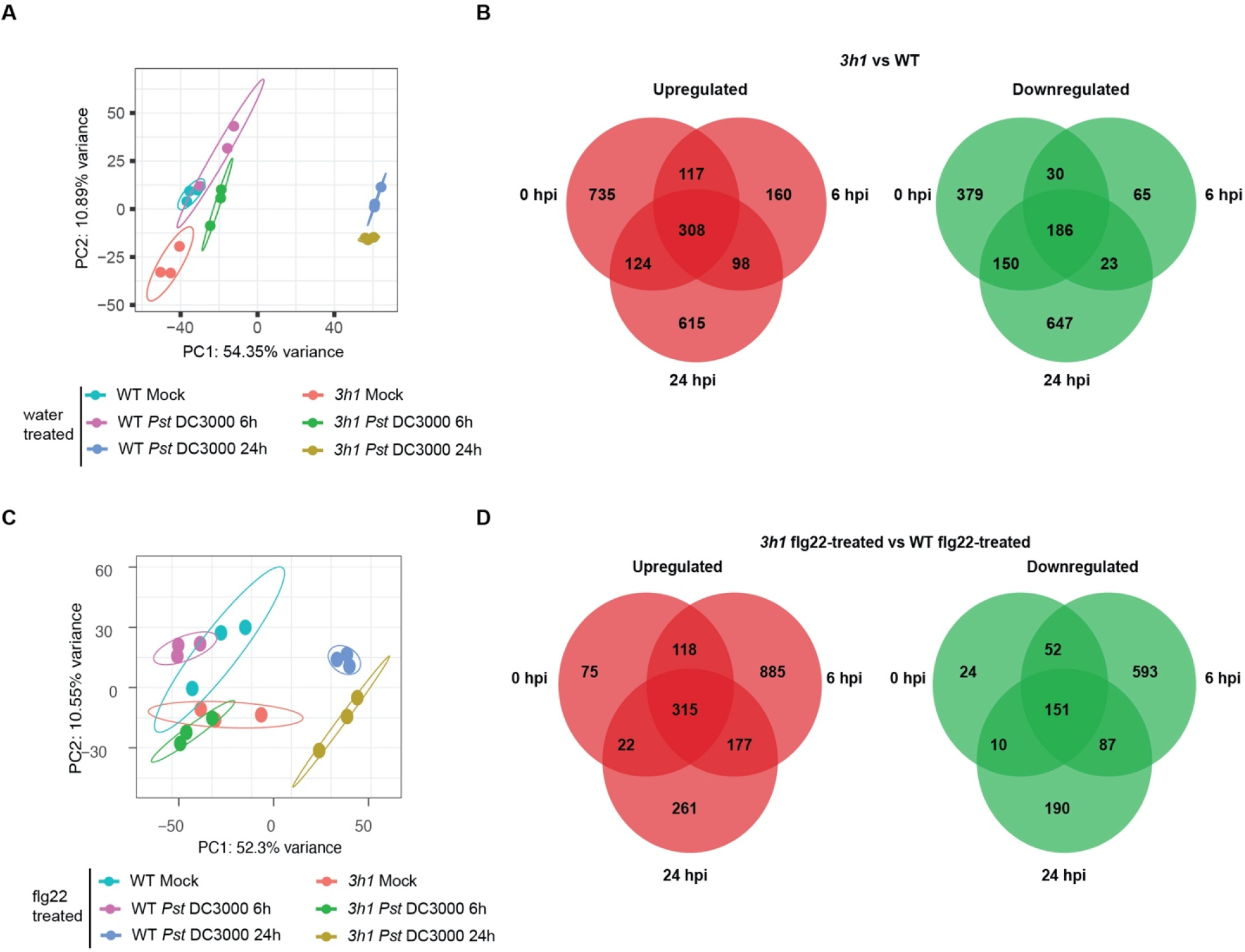
Transcriptome of *3h1* plants **(A)** Principal component analysis (PCA) plot of the RNA-seq of WT and 3h1 plants treated with *Pst* DC3000 infection at 6 and 24 h of infection. **(B)** Venn diagrams showing the number of upregulated and downregulated genes in *3h1* as compare to WT after *Pst* DC3000 infection. **(C)** Principal component analysis (PCA) plot of the transcriptome of flg22 pretreated WT and 3h1 plants challenged with *Pst* DC3000 infection at 6 and 24 h of infection. **(D)** Venn diagrams showing the number of upregulated and downregulated genes in flg22 treated *3h1* as compare to flg22 treated WT and then challenged by *Pst* DC3000.

**Figure EV3.**
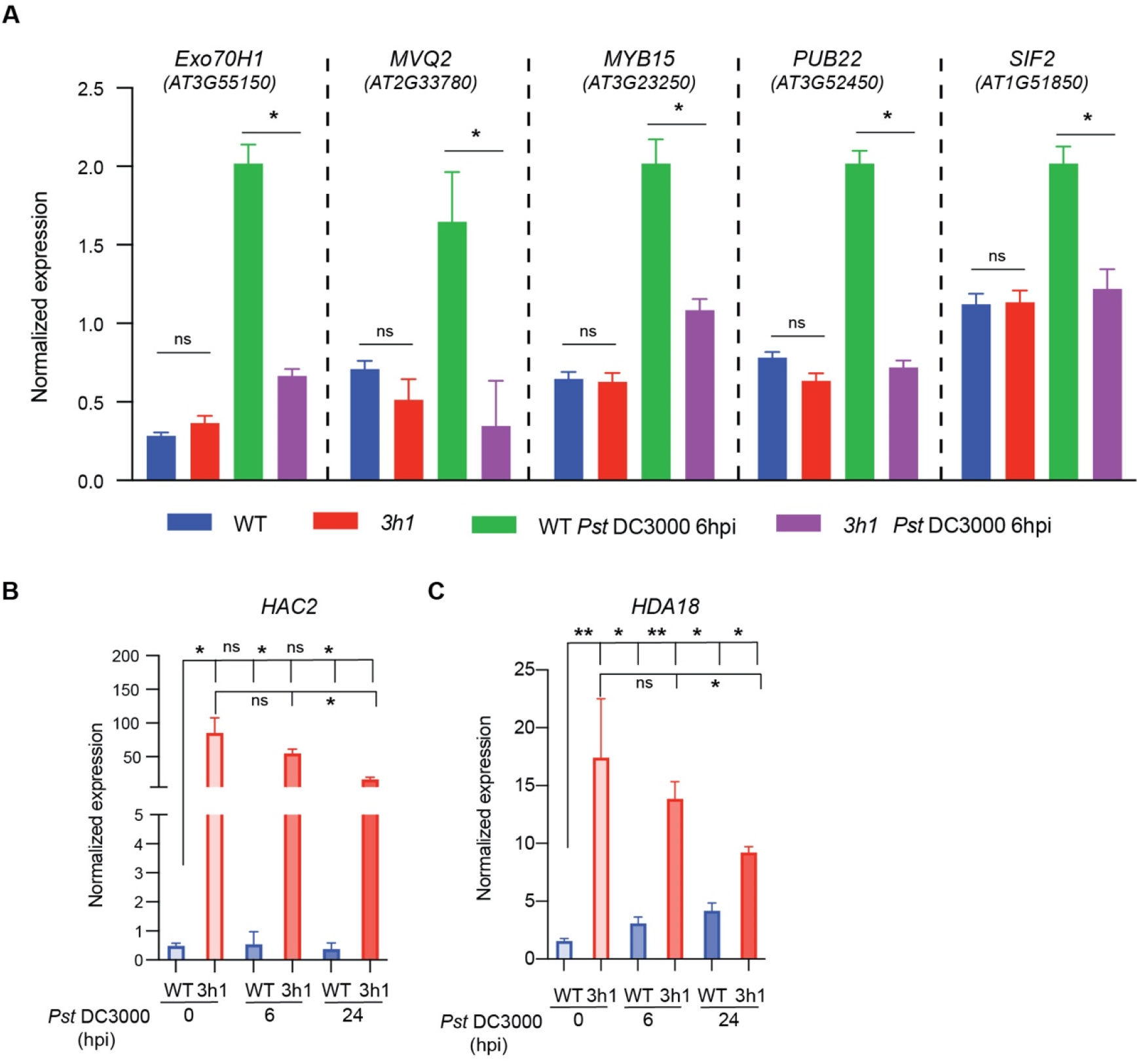
**(A)** qRT-PCR validation of indicated defense related genes which showed altered expression pattern between WT and *3h1* before and after infection when pre-treated with flg22. **(B)** qRT-PCR expression analysis of histone acetylation regulators *HAC2* and *HDA18* between WT and *3h1* before and after infection when pre-treated with flg22. *TUB6* was used for normalization.

**Figure EV4.**
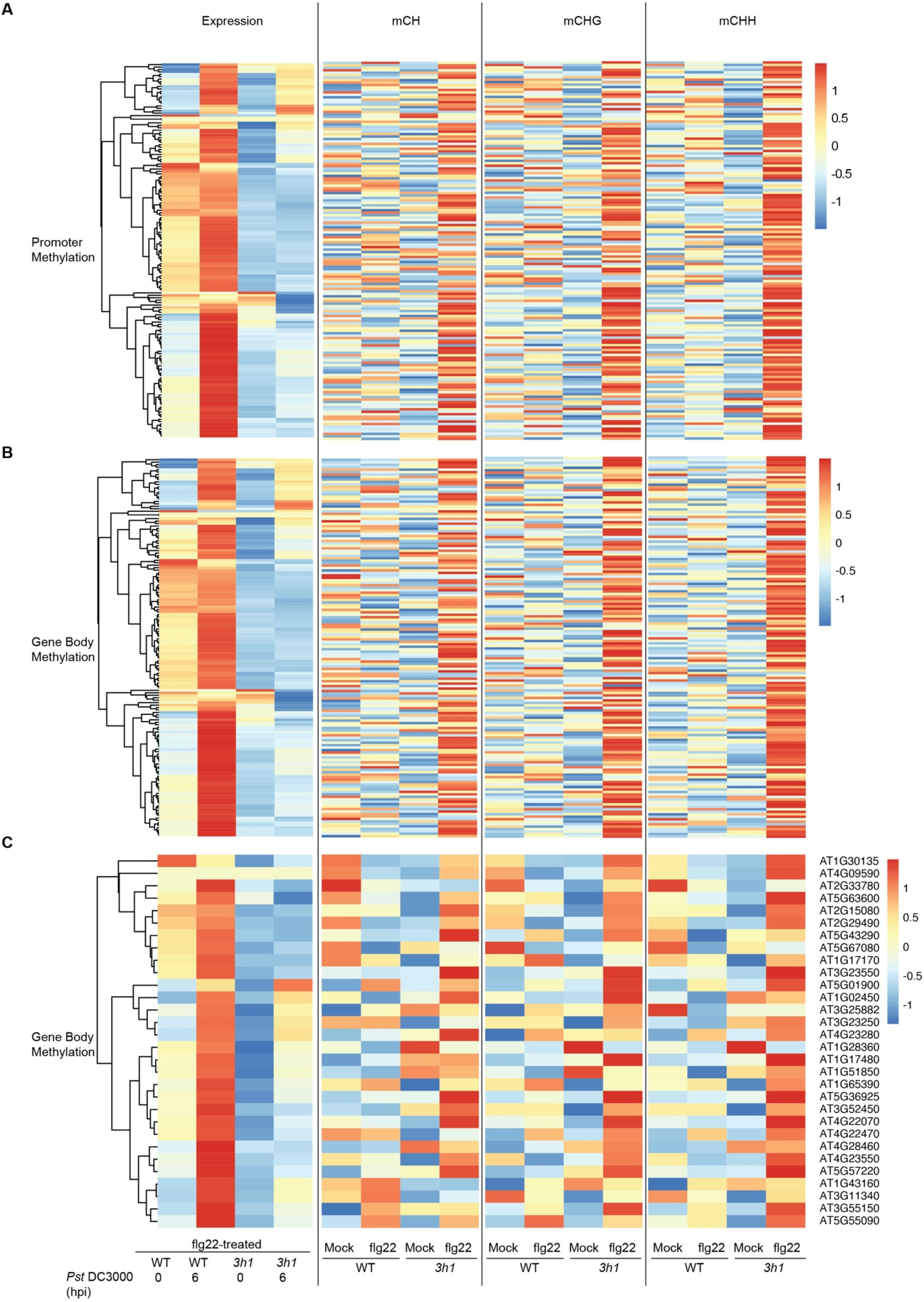
DNA methylation dynamics in *3h1* plants after flg22 treatment **(A, B)** Heatmap of mean promoter and gene body methylation in all contexts (CG, CHG and CHH) of Cluster 154 genes in Cluster 1 in flg22-treated WT and *3h1* samples which showed differential expression after *Pst* DC3000 infection (Fig 4D). **(C)** Heatmap of mean gene body methylation in all contexts (CG, CHG and CHH) on selected transcripts (30 genes) from Cluster 1 (154 genes) in flg22-treated WT and *3h1* samples which showed differential expression after *Pst* DC3000 infection (Cluster 1 of Fig 4D).

**Figure EV5.**
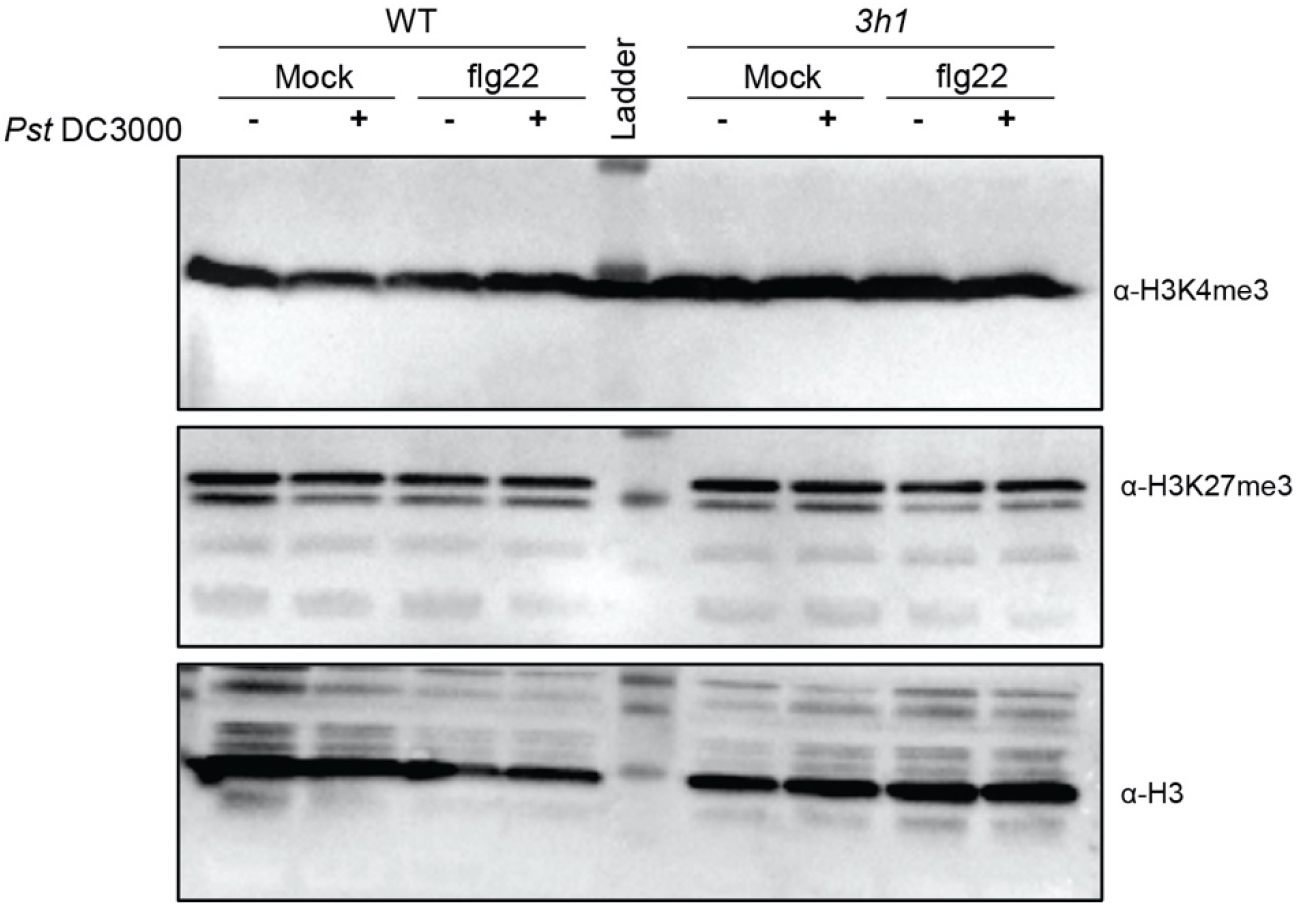
Western blot showing the H3K4me3 and H3K27me3 levels in WT and *3h1* plants after water (mock) or flg22 treatment challenged with *Pst* DC3000.

**Figure EV6.**
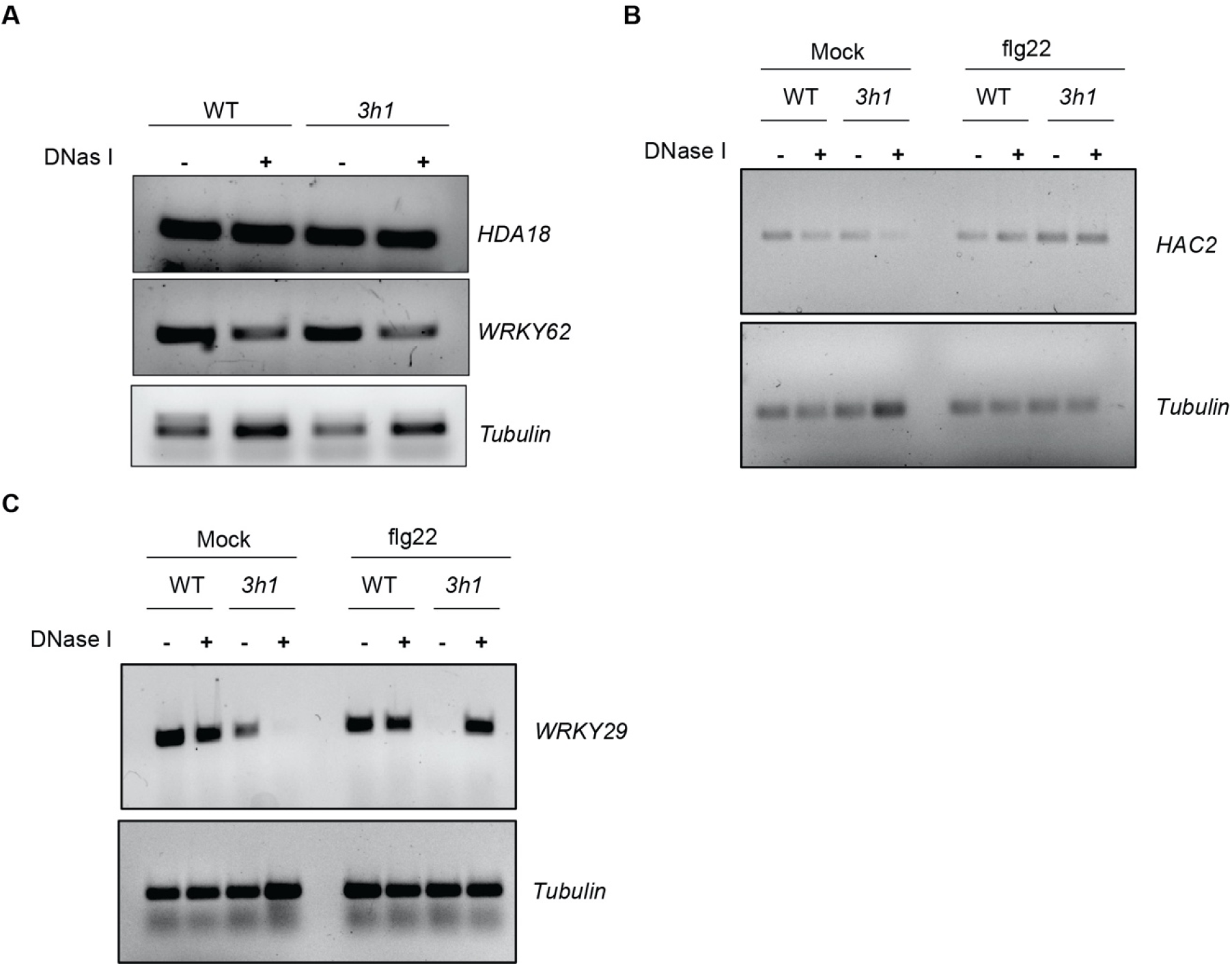
(**A**) DNase I accessibility PCR of *HDA18* and *WRKY62* in WT as compared to *3h1*. (**B, C**) DNase I accessibility PCR of *HAC2* and *WRKY29* in WT as compared to *3h1* before and after flg22 treatment.

## Acknowledgments

We are grateful to Dr. Kinga Rutowicz and Dr. Célia Baroux from DPMB, University of Zurich, Switzerland for kindly sharing the H1 related mutant seeds and fruitful discussions. We also thank KAUST core lab for providing RNA seq and hormone quantification facilitates.

## Author contributions

AHS and HH conceptualized this study. AHS performed most of the experiments. KN analyzed RNA seq and Bisulfite seq data. NT performed some of the qPCR experiments. MT did phytohormone quantifications. HA and NR helped in performing initial patho-assays. AHS, KN, NR and HH wrote the manuscript. All authors read and approved the final manuscript.

## Data and materials availability

RNA seq and Methylation data generated for this study have been deposited in the NCBI under accession number PRJNA814075. Additional data related to this paper may be requested from the authors.

---

**Table EV1.**
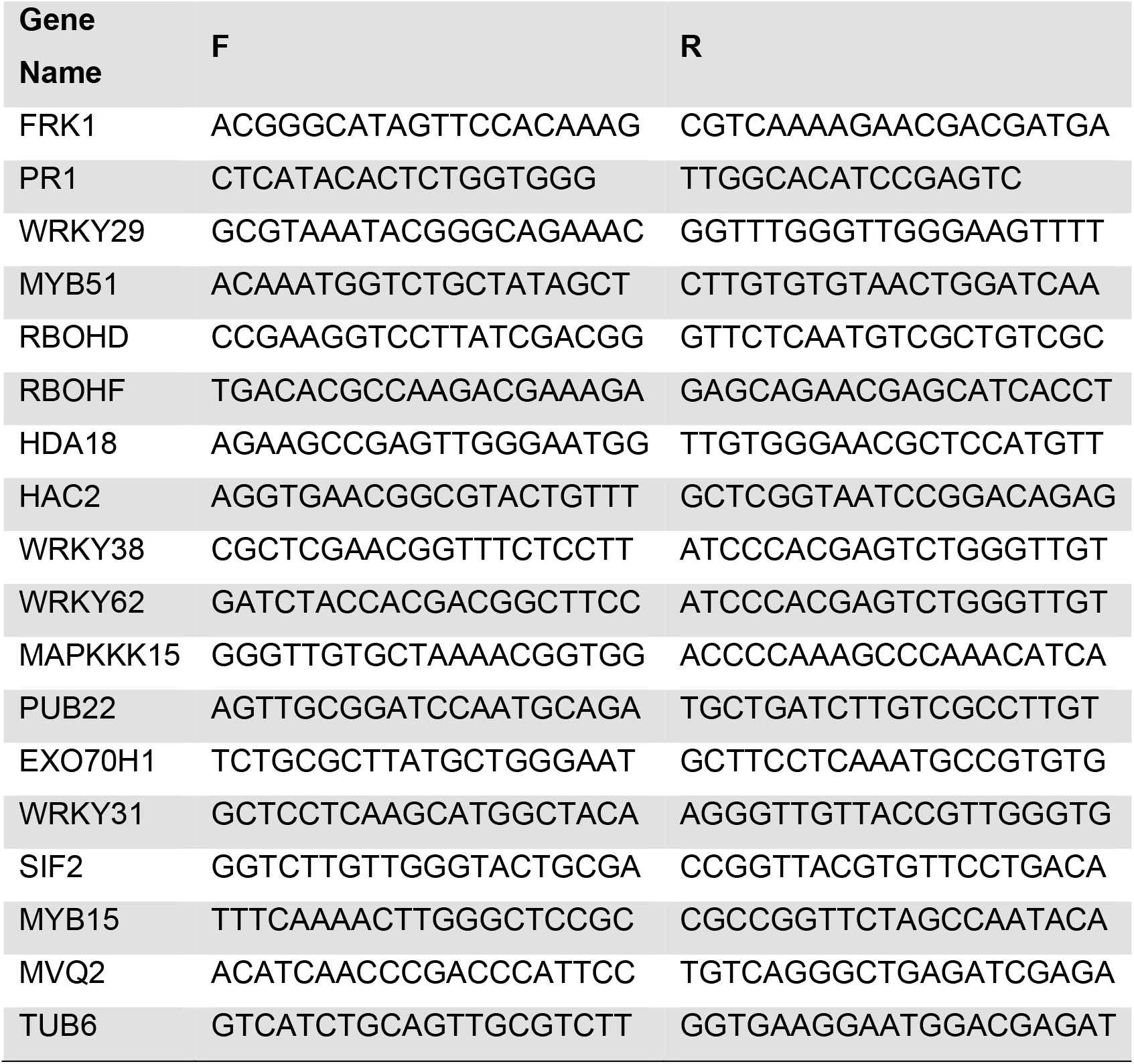

